# Structure-function properties in disordered condensates

**DOI:** 10.1101/2020.05.14.096388

**Authors:** Kamal Bhandari, Michael A. Cotten, Jonggul Kim, Michael K. Rosen, Jeremy D. Schmit

## Abstract

Biomolecular condensates appear throughout the cell serving a wide variety of functions. Many condensates appear to form by the assembly of multivalent molecules, which produce phase separated networks with liquid-like properties. These networks then recruit client molecules, with the total composition providing functionality. Here we use a model system of poly-SUMO and poly-SIM proteins to understand client-network interactions and find that the structure of the network plays a strong role in defining client recruitment, and thus functionality. The basic unit of assembly in this system is a zipper-like filament composed of alternating poly-SUMO and poly-SIM molecules. These filaments have defects of unsatisfied bonds that allow for both the formation of a 3D network and the recruitment of clients. The filamentous structure constrains the scaffold stoichiometries and the distribution of client recruitment sites that the network can accommodate. This results in a non-monotonic client binding response that can be tuned independently by the client valence and binding energy. These results show how the interactions within liquid states can be disordered yet still contain structural features that provide functionality to the condensate.

## I. INTRODUCTION

Many cellular structures have been shown to form by the spontaneous condensation of biomolecules into liquid-like states [1, 2], often through liquid-liquid phase transitions. While these condensates may contain hundreds of different molecules, typically only a small number of molecules with high interaction valence and high connectivity to other molecules in the structure contribute strongly to phase separation [3–5]. These are said to have scaffold-like properties depending on how strongly they promote phase separation. The remaining molecules, which exhibit client-like properties, are recruited through interactions with scaffolds [6, 7]. Together, the collection of molecules in a condensate determines its functionality.

Since the molecules driving phase separation are multivalent, polymer-like species, many treatments of condensate formation are based on polymer theories [8–11]. In particular, scaffold condensation can be understood as the interaction between attractive stickers separated by inert spacers [11, 12]. These efforts explain universal features of condensates, such as how multivalency can amplify the effect of weak interactions to tune the phase coexistence line to lie within physiologically relevant regions of phase space (e.g. physiological concentrations) [8]. However, the diverse functions performed by different condensates requires that there are also non-universal features [2, 7, 13–16], and each of these assemblies will be under evolutionary pressure to optimize its specific function. This begs the question of how the disordered network of interactions within a liquid structure can affect its properties.

The disorder in biomolecular condensates poses a challenge for structural biology to determine what features of the assembly are functionally relevant. In previous work, we showed that analytic theory can be used to deduce structural features in these systems [17]. Here we apply this approach to condensates formed by poly-SUMO and poly-SIM, a synthetic system with a 1:1 binding stoichiometry between SUMO and SIM modules that was developed to study the recruitment of client molecules [6]. This system captures the features of the common sticker-and-spacer topology, without the ambiguity in identifying the sticker moieties that complicates the study of natural systems. We find that SUMO/SIM condensates have a microstructure that differs from other condensates [11, 17]. This microstructure dictates how client recruitment varies with changes in valence and affinity, and suggests that the composition, and thus function, of a condensate can be changed through modulation of its internal structure independent of its propensity to undergo phase separation. These results show how functionally relevant structures can be embedded within liquid-like disorder.

We propose a general model in which functional properties emerge from hierarchical assembly within a condensate. In this model, strong interactions result in the formation of molecular complexes with functional properties. These complexes then condense into a fluid state via weaker interactions. This combination of strong and weak interactions combines the best features of both interactions modes. Strong interactions provide the structural specificity needed for functional properties to emerge, while weaker interactions allow the formation of a high density state without the kinetic arrest that would accompany assembly driven strictly by strong interactions.

## RESULTS

### Zipper-like filaments allow most modules to form bonds within a sparse network

Our initial attempts to model SUMO/SIM condensates using three-component Flory-Huggins theory or a modified version of the model recently applied to SPOP/DAXX assemblies [17] yielded wildly inaccurate predictions for the previously reported condensate density [6]. The high density predicted by these models can be understood with a simple dimensional analysis argument. Within a random network, we expect the sticker density to be on the order of 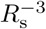, where *R*_s_ is the radius of gyration of a sticker+spacer repeat unit. In the poly-SUMO/poly-SIM system *R*_s_ *∼* 3.3 nm, which leads to a droplet module concentration of 50 mM, much greater than the observed concentration of *∼* 2 mM [6]. The latter concentration implies that the volume per decavalent scaffold is (20 nm)^3^, which would allow only a small fraction of intermolecular bonds to be satisfied if the molecules are randomly oriented. For bond energies strong enough to drive phase separation, on the order of *k*_*B*_*T*, the penalty for this many unsatisfied bonds is prohibitive.

These considerations require that SUMO/SIM networks satisfy most binding sites while simultaneously maintaining an average molecular spacing on the order of the scaffold dimension. These requirements can only be fulfilled if the molecules are aligned to form zipper-like structures (Fig. 1a). These zippers will have defects in the bonding structure that allow for the subsequent formation of a 3D network. Fig. 1a shows two defects considered in this work, gaps and sticky ends. Other defects such as cyclization and overlaps, are also possible but neglected in the current treatment. These defects provide recruitment sites for clients as well as enable the 1D filaments to assemble into a 3D network.

**FIG. 1:**
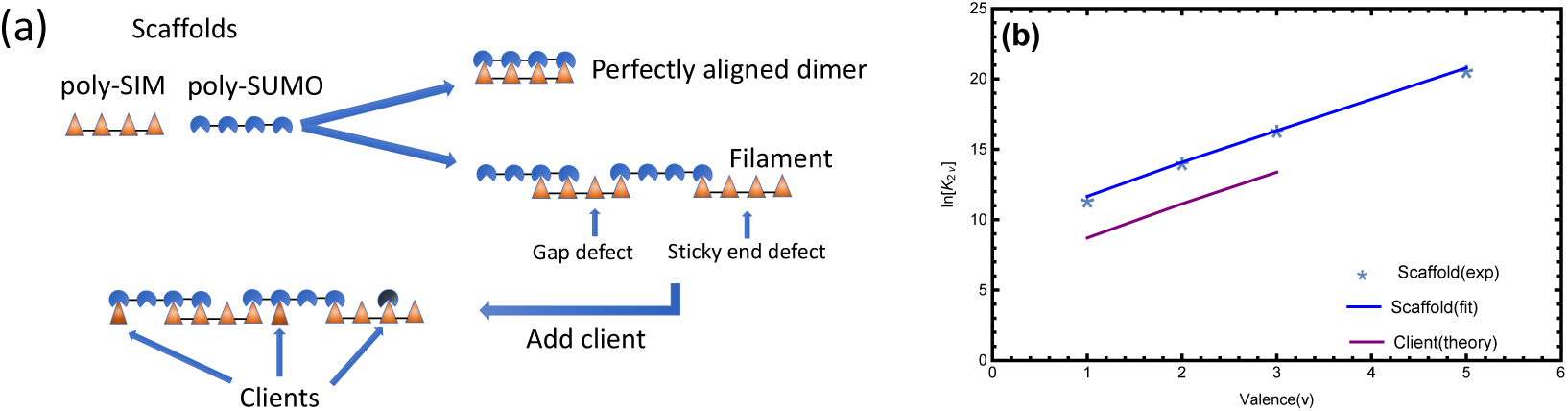
(a) Poly-SUMO and poly-SIM assemble into zipper-like filaments. The short linker connecting modules favors consecutive bonds with the same molecule rather than the formation of a random network. The filaments contain defects of unpaired modules in the interior of the filament (“gaps”) and at the ends (“sticky ends”). These unsatisfied bonds allow for the crosslinking of filaments into a 3D network and the binding of clients (bottom row). (b) The dimerization of low valence poly-SUMO/poly-SIM constructs provides model parameters. Plot of the dimer association constant, *K*_2*v*_ [6], vs. the valence, *v*. The module binding free energy, *E*, is obtained from the slope of ln *K*_2*v*_ (blue line), while the intercept provides the reference concentration *c*_0_. Clients (mono-, bi-, or trivalent SIM) have a lower binding affinity than scaffolds in the droplet phase, which we attribute to steric interactions between the network and the fluorescent labels. We account for this with a free energy offset, *f*_RFP_, for clients in the dense phase (purple line).

### The filament partition function can be calculated with transfer matrices

Due to the 1D nature of the filaments, the defect density can be conveniently calculated using a transfer matrix formalism [18, 19]. To illustrate this approach, consider a tetravalent (*v* = 4) system (in subsequent calculations we use *v* = 10 to compare to the experiments of [6]) and define *Q*_*N*_ (*j*) as the partition function of a filament containing *N* molecules that terminates with a sticky end of *j* unpaired sites. Next, we construct a *v −* 1 dimensional vector, **V**(*N*), whose elements are the statistical weights *Q*_*N*_ (*j*) for sticky ends of each length, *j*. This vector obeys a recursion relation **V**(*N* +1) = **M**_*i*_**V**(*N*), where the transfer matrix **M**_*i*_ generates the possible states upon addition of a molecule to the filament

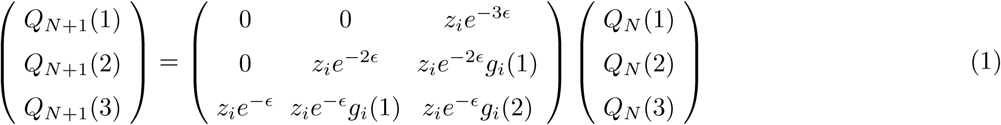

where *ϵ* is the SUMO-SIM binding free energy (all energies are expressed in units of *k*_*B*_*T*) and *z*_*i*_ is the fugacity of the added molecule. Matrix elements below the antidiagonal leave gap defects which can bind clients or form crosslinks with other filaments. This is accounted for by the gap partition function *g*_*i*_(*m*), where *m* is the size of the gap. Since we have two scaffold molecules, the grand partition function for a filament of *N* + 1 poly-SUMO and *N* + 1 poly-SIM molecules can be generated by alternately applying the matrices for the two species and applying vectors **V**_L*/*R_ to collapse the matrix product into a scalar polynomial (see Methods). This operation is given by *Q*_*N*+1_ = **V**_L_(**M**_U_**M**_I_)^*N*^ **V**_R_, where the U/I subscripts denote SUMO/SIM.

The most important parameter in Eq. 1 is the module binding affinity *ϵ*, which can be obtained from dilute solution measurements [6] as follows. Consider the dimer association constant between scaffolds of valence *v*, 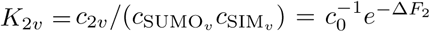, where *c*_2*v*_ is the concentration of scaffold dimer. Here the reference concentration *c*_0_ gives the equilibrium constant appropriate units and enters the matrix formalism through the fugacity *z*_*i*_ = *c*_*i*_*/c*_0_ (valid for dilute solution when the monomer free energy is set to zero). The dimerization free energy is given by the dimer partition function 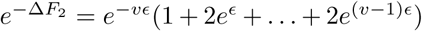, where the terms represent the perfectly aligned state and successively larger mis-alignments. Plotting ln *K*_2*v*_ vs. *v* (Fig. 1b), we can extract *ϵ* = *−*2.23 *k*_*B*_*T* from the slope and *c*_0_ = 64 *µ*M from the intercept.

To compute solution properties we construct the grand partition function

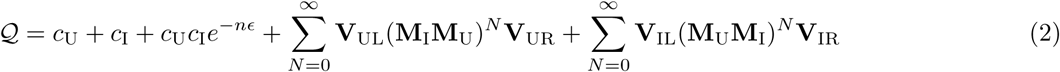

where the first two terms represent the monomers, the third term represents the perfectly aligned dimer, and the two sums represent filaments starting with SUMO and SIM respectively. The matrices **M**_*i*_ and end vectors **V** are given in the Methods section.

### Theory parameters describe fluid structure

The zipper assemblies give the fluid a hierarchical structure. At short length scales the condensate is characterized by zippers stabilized by cooperative binding, while at longer length scales it is characterized by a 3D network stabilized by monovalent crosslinks. While the weak crosslinks provide the system with fluidity, here we focus on two parameters that describe the short length scale structure that provides functionality. The first is the defect density, which describes the capacity of the filaments to bind clients or crosslink into a 3D network. Using the substitution *g*_*i*_(*m*) = *g*^*m*^ so that *g* is the statistical weight of an unbound module, the density of unpaired sites per scaffold molecule is given by 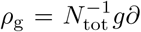. This gives *ρ*_g_ ≃ 0.6 unbound sites per decavalent scaffold at equimolar scaffold mixtures (Fig. 2a).

**FIG. 2:**
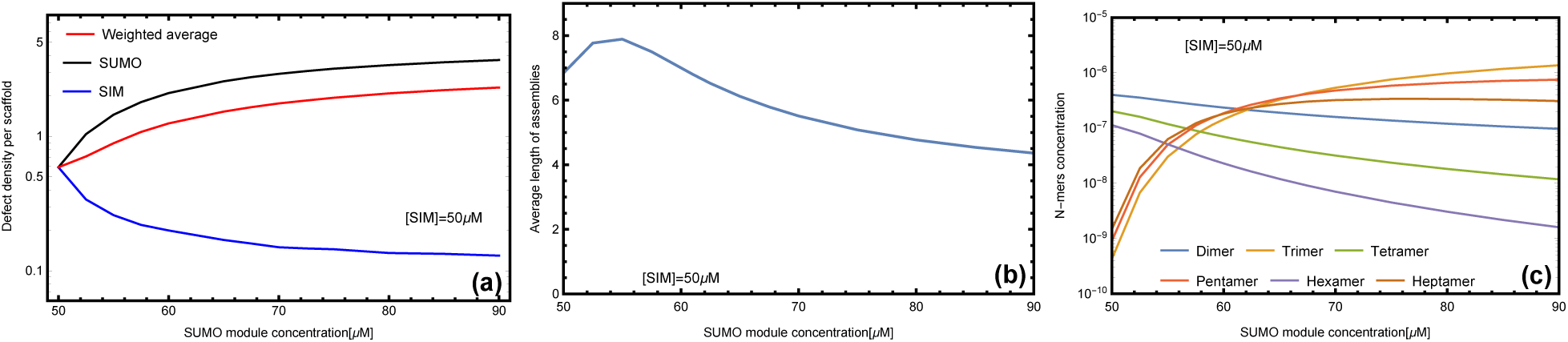
Filament length and defect density depend on stoichiometry. (a) At equal stoichiometry most scaffolds are fully bound resulting in few defects. As the stoichiometric imbalance increases, the number of unbound SUMO modules increases while the number of unbound SIM sites decreases. (b) The average length of filaments is a non-monotonic function of scaffold stoichiometry. A small excess of one scaffold leads to an increase in the filament length because unpaired scaffolds are available to stabilize sticky ends resulting from mis-aligned states. Larger stoichiometric mismatches result in a decline in filament length, which provides more sticky ends to bind the excess scaffold. (c) Filaments with equal number of SUMO and SIM scaffolds are favored at symmetric mixing, but unequal stoichiometries favor odd length filaments.

The second useful parameter is the average number of scaffolds in a filament. This quantity is obtained from *⟨N*_i_*⟩* = *z*_*i*_*∂* ln *𝒬/∂z*_*i*_. The total assembly size *N*_tot_ = *⟨N*_I_ + *N*_U_*⟩* is plotted in Fig. 2b. The most energetically favored state is the perfectly aligned dimer which allows all binding sites to be satisfied (Fig. 1a). This competes with the higher entropy of misaligned states, particularly upon the formation of filaments with *N*_tot_ *≥* 3 because the unpaired sites can be distributed throughout the filament rather than localized at the ends. At sufficiently high concentration the entropic gain can overcome the energetic cost of misalignment defects allowing the formation of large assemblies.

### Asymmetric scaffold stoichiometries promote filaments over perfectly aligned dimers

The combined behavior of the defect density and the filament length can be understood by looking at the populations of different filament lengths (Fig. 2c). At equal stoichiometries, the solution is dominated by perfectly aligned dimers resulting in a small average complex size. This is similar to the “magic number” effect seen in the simulations of EPYC1 and Rubisco [20, 21]. At unequal stoichiometries the energetic penalty for misalignments is reduced because excess scaffolds are available to bind one of the sticky ends, which facilitates the formation of longer complexes. As the stoichiometry mismatch increases, the system becomes dominated by odd-length complexes which are initially large before trimer (2:1 stoichiometry) states dominate at large stiochiometric mismatches. The decrease in filament size at large stoichiometric asymmetry can be understood as an abundance of filaments that have two sticky ends of the same type, which prevents them from joining into larger assemblies. The shortening of filaments causes an increase in sticky ends and, therefore, an increase in the concentration of defect sites (Fig. 2a). The defect density increases rapidly at first, then slows at SUMO module concentrations above 70 *µ*M as the droplet becomes dominated by the 2:1 trimer.

### Asymmetry in scaffold stoichiometry can only be accommodated at the filament ends

Our matrix formalism does not allow for direct calculation of the assembly of filaments into a 3D network within the droplet. However, we can compute the ratio of SUMO to SIM scaffold in the droplet by assuming that the monomer and perfectly aligned dimer remain in the bulk solution, *𝒬*_dilute_ = *c*_U_ + *c*_I_ + *c*_U_*c*_I_*e*^*−vϵ*^, while all other complexes are in the droplet, *𝒬*_droplet_ = *𝒬 −𝒬*_dilute_. The calculated stoichiometry is compared to the experimentally measured droplet composition in Fig. 3a. The droplet SUMO/SIM ratio has downward curvature indicating that the droplet is unable to accommodate the stoichiometry mismatch of the bulk solution. This is a consequence of the filamentous structure because the interior of the filament is constrained to a 1:1 stoichiometry, so stoichiometric imbalance can only be accommodated at the filament ends. The filament calculation has somewhat more curvature than the experimental ratio (Fig. 3a, solid line) suggesting that the approximation of purely 1D assembly breaks down when the SUMO/SIM ratio is greater than *∼* 1.4. Under these conditions, the concentration of free scaffold (Fig. 3b) is sufficiently high that scaffold binding to gap defects cannot be neglected. A simple correction, allowing monovalent binding to the gap defects (see Methods), resolves the discrepancy (Fig. 3a, dashed line).

**FIG. 3:**
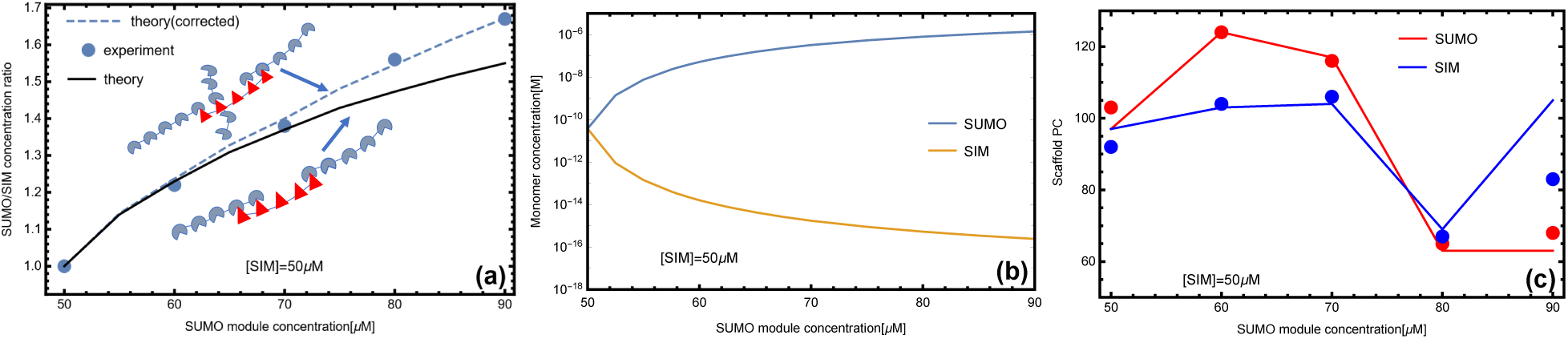
Zipper structure constrains droplet stoichiometry. (a) The ratio of poly-SUMO to poly-SIM (*N*_U_*/N*_I_) calculated from our theory agrees well with the droplet stoichiometry in experiments [6]. At high stoichiometric mismatches the approximation of purely 1D filaments breaks down because the concentration of free scaffolds is high enough to allow binding in the gaps. A correction accounting for monovalent scaffold-gap binding (inset cartoon) resolves the discrepancy with experiment (dotted line). (b) Excess SUMO scaffold accumulates in the dilute phase and depletes the concentration of monomeric SIM scaffolds. (c) Small stoichiometric mismatches promote increased scaffold accumulation in the droplet phase, but the trend reverses as the average filament size drops. The discrepancy at 90 *µ*M can be explained by the “t” configuration in the cartoon of panel (a).

A related calculation is to compare the scaffold partition coefficient (PC), defined as the concentration in the dense phase to the concentration in the dilute phase. The fractional composition in each phase is *N*_*i*_*/*(*N*_U_ + *N*_I_), where *i* = U or I. In order to compare with experiments we weight this quantity by the experimentally determined total module concentration in each phase. Fig. 3c plots the quantities

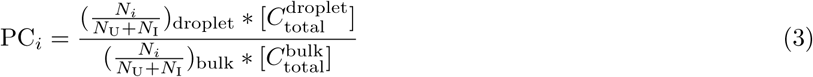

Where 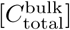 and 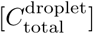 are the total module concentration in bulk and droplet phase respectively [6]. The initial rise in the scaffold partitioning is due to the increase in filament lengths, but the trend reverses with the accumulation of smaller filaments at large stoichiometric asymmetry.

### Clients can bind to filament defects

We next examined how the structural properties of scaffold assemblies in the droplet and bulk phases influence partitioning of mono-, di-, and trivalent SIM molecules employed as clients by Banani et al. [6]. In particular, we sought to understand the curious observation that high-valence clients show non-monotonic partitioning behavior as the SUMO/SIM scaffold ratio changes [6]. To account for the recruitment of clients to the droplet we add client binding to the gap partition function. The form of the gap partition function depends on the valence and the concentration, *c*_cl_, of the clients as follows

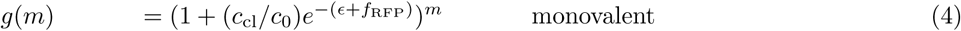

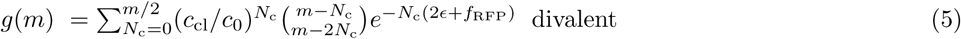

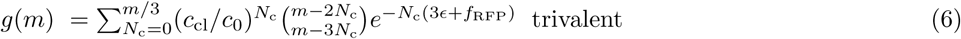

In the case of monovalent clients, each gap site is independent, which gives a simple binomial with the two terms representing the empty and client-bound states. For multivalent clients we have the more involved process of counting the number of ways to add *N*_c_ clients to *m* sites and summing over the number of clients that will fit in a gap (see Methods). While the binding energies in the above expression are same as those for the scaffolds due to the same underlying SUMO-SIM interaction, the binding affinity of clients is lower in the droplet than in the bulk [6]. We attribute this to the entropy cost of confining the bulky RFP fluorescent tag within the droplet network. This steric penalty is included in our model with the factor *f*_RFP_. A least squares fit of the client partitioning yields *f*_RFP_ = +3.15 *k*_*B*_*T* (Fig. 1b). *f*_RFP_ is the only free parameter in our model, and has the effect of altering the peak hight with minimal effect on the peak location (Fig. 5).

Accounting for bound and unbound client in each phase, the client partition coefficient can be expressed as

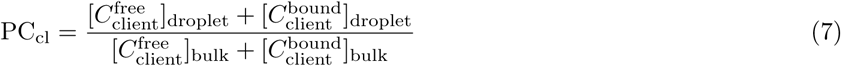

where 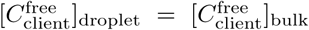 are the free client concentrations in the droplet and bulk, respectively, and 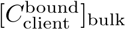 and 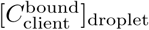 are the bound client concentration in the bulk and droplet phase, respectively. The average number of bound clients per scaffold is computed as 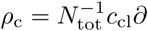. The bound client concentration is obtained by multiplying *ρ*_c_ times the experimentally determined scaffold concentration [6]. Similarly, the bound client concentration in the dilute phase 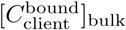 can be calculated from the total number of clients per scaffold in the dilute phase multiplied by the corresponding scaffold monomer concentration in the bulk phase.

Note that the concentrations *c*_U_, *c*_I_, and *c*_cl_ appearing in the partition functions all represent the concentration of molecules that have not formed intermolecular bonds. These values must be determined by numerically solving the equations *C*_tot_ = *c*_*i*_*∂𝒬/∂c*_*i*_ for scaffolds and 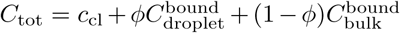 for the client. Here *C*_tot_ is the total (bound and unbound) concentration of the relevant molecule and *φ* is the volume fraction occupied by the droplet phase. The droplet volume fraction can be determined from the experimental values [6] of the dilute and droplet protein concentrations using *C*_tot_ = *φC*_droplet_ + (1 *− φ*)*C*_dilute_.

### Filament defects compete with soluble scaffold molecules for client binding

The most striking feature of the PC is the non-monotonic dependence on the poly-SUMO concentration (Fig. 4a). This can now be understood as follows. Small excesses of poly-SUMO scaffold are readily incorporated in the droplet, which increases the PC by providing more sites for SIM clients to bind. However, as previously discussed, the 1D filaments are limited in their ability to accommodate unequal stoichiometry. As the filament ends become saturated with the abundant scaffold species, excess scaffold monomers are forced to accumulate in the dilute phase (Fig. 3b). At SUMO concentrations above 75 *µ*M the number of unpaired SUMO modules in the dilute phase exceeds those in the dense phase (Fig. 4b). These free scaffolds compete for clients, which increases client concentration in the dilute phase and lowers the PC.

**FIG. 4:**
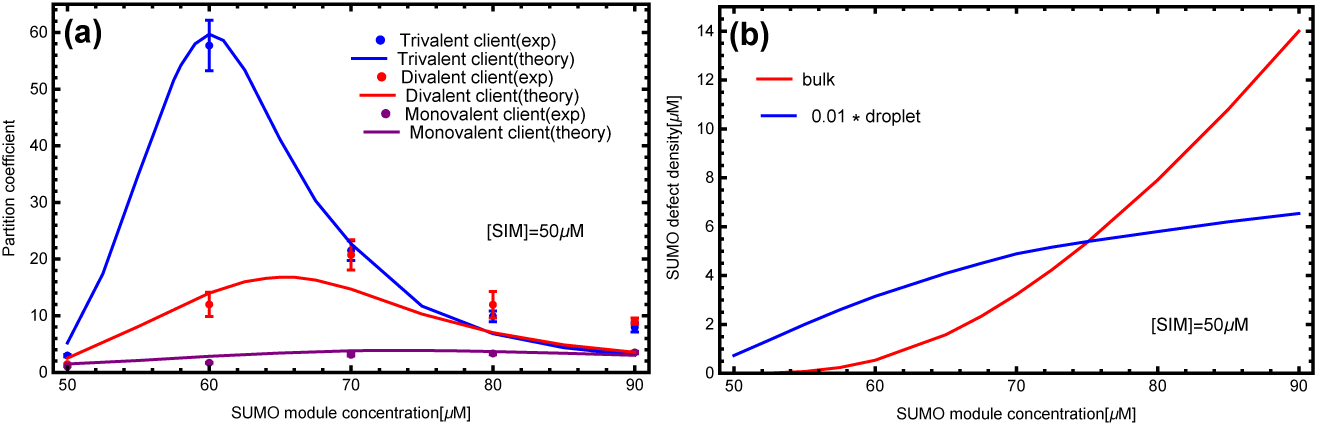
The scaffold composition that optimizes client recruitment depends on the client valence. (a) The transfer matrix theory (lines) captures the shift in partition coefficient peak as the client valence increases. The high affinity of trivalent clients is more sensitive to both the appearance of defects in the droplet and the presence of free scaffold in the bulk. Experimental data from [6]. (b) Concentration of unbound SUMO modules in the dense and dilute phases. Excess scaffolds, and associated defect sites, are initially bound in the droplet as shown by the blue curve (given by *ρ*_*g*_ times the droplet poly-SUMO concentration). However, above 70 *µ*M additional poly-SUMO accumulates primarily in the bulk phase, which has a defect density of 10*c*_U_ (red). The scaling factor applied to the droplet curve is approximately equal to the *∼* 0.01 volume fraction of the droplets. Therefore, these curves approximately represent the number of defects sites in each phase.

**FIG. 5:**
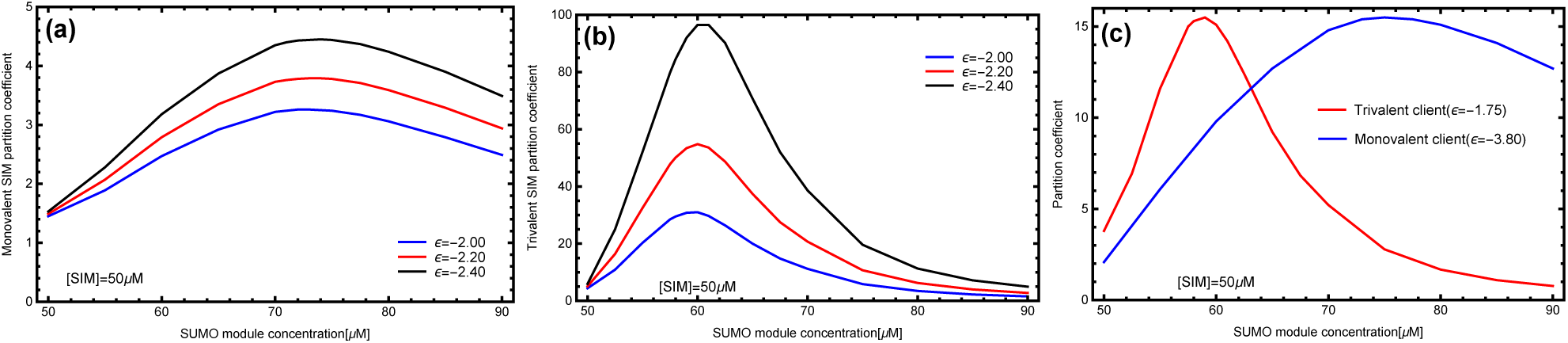
Increasing the client module binding affinity enhances client recruitment with minimal effect on the location of the peak. This provides separate mechanisms to tune the location and magnitude of client recruitment. (a) Monovalent SIM and (b) Trivalent SIM at different module binding affinities. The different effects of client valence and affinity on the PC curve allow the network to switch between the recruitment of different clients. (c) With tuned client affinities, it is possible for the network to selectively recruit either the monovalent or trivalent client in different regimes of parameter space.

The client binding response is sensitive to the valence and binding affinity of the clients. Increasing the client valence increases the maximum value of the PC and shifts the peak toward equal scaffold stoichiometry. The increased affinity of clients through higher valence allows them to bind more readily to the sticky ends that appear with a small excess of complementary scaffold. However, the larger valence of these clients means that there is comparatively lower configuration entropy when binding to small defects. Therefore, when free scaffolds appear in the dilute phase, the favorable entropy provided by 10 consecutive binding sites overwhelms gap binding. Conversely, low valence clients have lower affinity, requiring a higher defect density for appreciable recruitment. But, they do not feel a confinement entropy in small defects, so they are sensitive only to the total number of available binding sites in the two phases (Fig. 4).

### Network structure can provide binding specificity

Not surprisingly, increasing the client’s specific affinity enhances client recruitment and increases the PC (Fig. 5a,b). Therefore, the filament microstructure of the droplet provides independent mechanisms to tune the magnitude (via the affinity) and concentration (via the valence) of the maximial PC. These orthogonal methods of tuning client recruitment provide a mechanism by which the structure of the network influences the recruitment of clients, and thus function of the droplets. Fig. 5c shows PC curves for a monovalent and trivalent client where the binding affinities have been tuned to have similar peak recruitment. However, because the peaks in the PC curves occur at different scaffold stoichiometries, the network will predominantly recruit either one client or other at different scaffold ratios. This means that, despite the liquid characteristics of the condensate, there is enough structure to specifically select between two closely related clients. Specifically, the zipper structure produces correlations in the location of available binding sites that biases the ensemble of binding states for multivalent clients.

## DISCUSSION

### Interactions stabilizing liquid states impart functionally relevant condensate properties

Comparing these results to the recently determined microstructure of SPOP/DAXX condensates [17], an interesting trend emerges. Both the SUMO/SIM and SPOP/DAXX systems can be described as sticker and spacer motifs [22, 23], yet the underlying networks have very different connectivity (Fig. 6). Despite the superficial differences, both systems assemble in a hierarchical manner; strong interactions form molecular complexes and weaker interactions drive the condensation of the complexes. In the SPOP/DAXX system, strong SPOP-DAXX interactions result in the formation of brush-like assemblies that condense into a liquid phase via weaker DAXX-DAXX interactions. However, at higher SPOP-DAXX ratios, DAXX crosslinks SPOP into rigid bundles that associate into a gel state. These distinct states allow the scaffold stoichiometry to serve as a kinetic switch between fluid and arrested states [17]. In contrast, in the poly-SUMO/poly-SIM system the same SUMO-SIM interactions drive the strong filament formation and the weaker network crosslinks. Also, the filamentous structure, which is favored by a spacer that is much shorter than the mean intermolecular spacing, allows for a sensitive control over client recruitment. In both systems the assemblies are disordered at the length scales of both the strong and weak interactions. However, the hierarchy gives structure to the fluid in a way that provides functional properties.

**FIG. 6:**
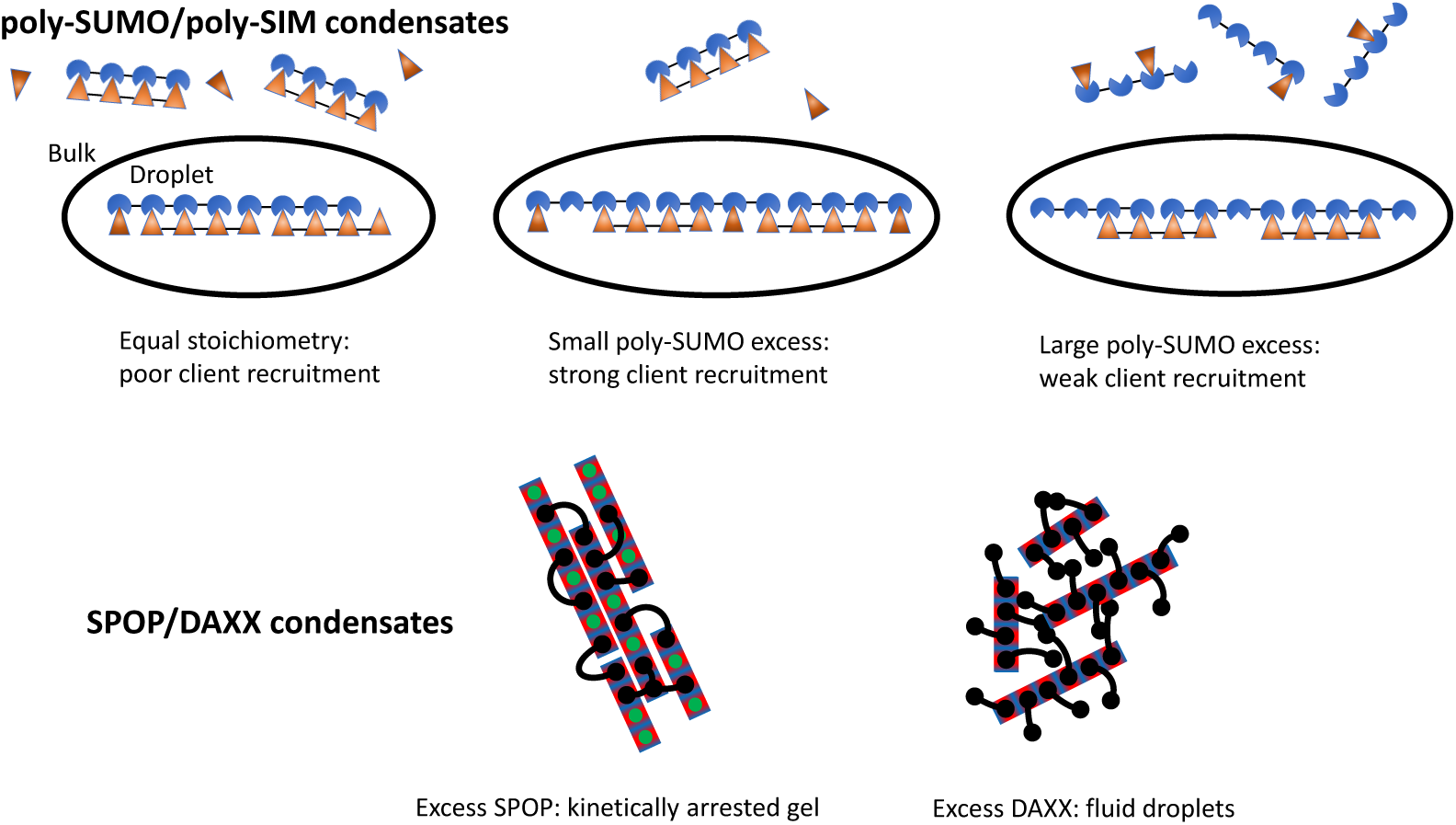
The microscopic connectivity of biomolecule condensates imparts specific properties. While both poly-SUMO/poly-SIM and SPOP/DAXX condensates form by the association of multivalent molecules, the assemblies have very different properties. Poly-SUMO/poly-SIM condensates are composed of linear filaments that provide a client binding response that is sensitive to the scaffold stoichiometry. SPOP/DAXX assemblies contain a “kinetic switch” that allows the system to convert between gel states with arrested dynamics and fluid droplets [17]. Here the black lines represent bivalent DAXX molecules, while the rectangles represent polymerized SPOP rods. Adapted with permission from [17]. Copyright 2020 by the American Chemical Society.

We conclude that even within a conserved sticker and spacer framework, differences in the scaffold valence, linker length, rigidity etc. can result in networks with widely different properties. It is inevitable that evolution will exploit these differences to optimize each condensate for its specific biological function. This inspires the question of how to identify functionally relevant structure within networks that are visually disordered. Here we have shown that theoretical modeling is a powerful tool for this task because parameters like *N*_tot_ and *ρ*_g_ emerge naturally to describe structural features in a way that connects to droplet function.

## IV. METHODS

### A. End Vectors and Transfer Matrices

Total grand partition function (Eq. 2) of the system is:

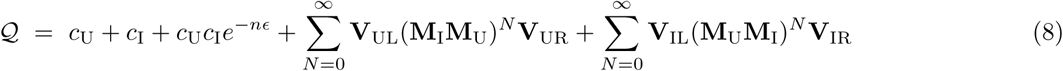

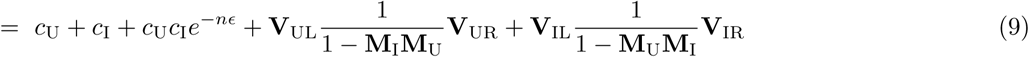

The two summation terms represent filaments starting with SUMO or SIM modules. In our formalism, filaments begin at the right end and grow right to left by application of the transfer matrices. The starting vectors **V**_UR_ and **V**_IR_ initiate the assembly with an imperfectly aligned dimer. These dimers have *v −* 1 different states where *v* is the valence of the scaffolds. These states have free energy *e*^*−*(*v−n*)ϵ*ϵ*^*g*_*i*_(*n*) where *n* = 1, 2, 3 *v −* 1, and *g*_*i*_(*n*) is the gap partition function for the sticky end defect at right end of the filament. These alignment states comprise the elements of the right vectors

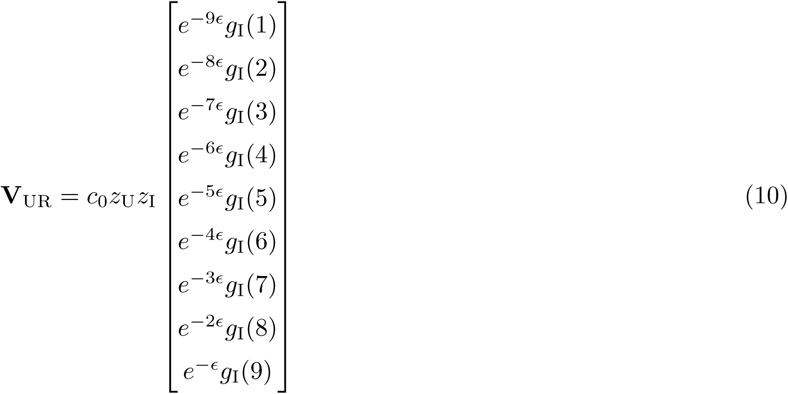

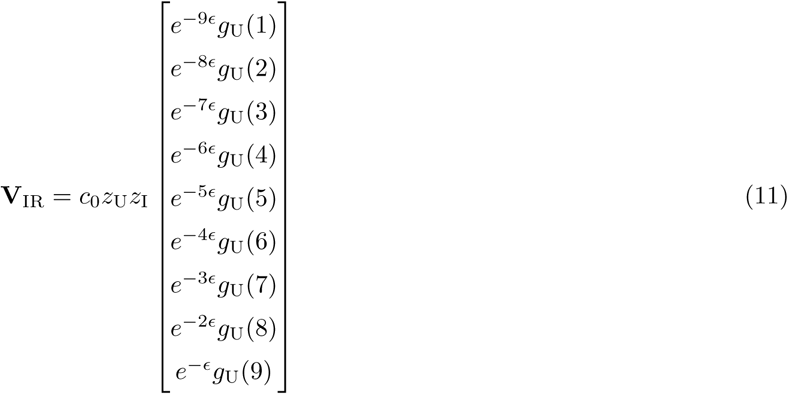

**V**_UR_ and **V**_IR_ are identical apart from the client species that can bind in the gap partition functions *g*_U_ and *g*_I_, where the gap partition functions are labeled with subscripts that describe the client that they bind.

The transfer matrix **M**_**i**_ for deccavalent (*v* = 10) scaffolds is given by

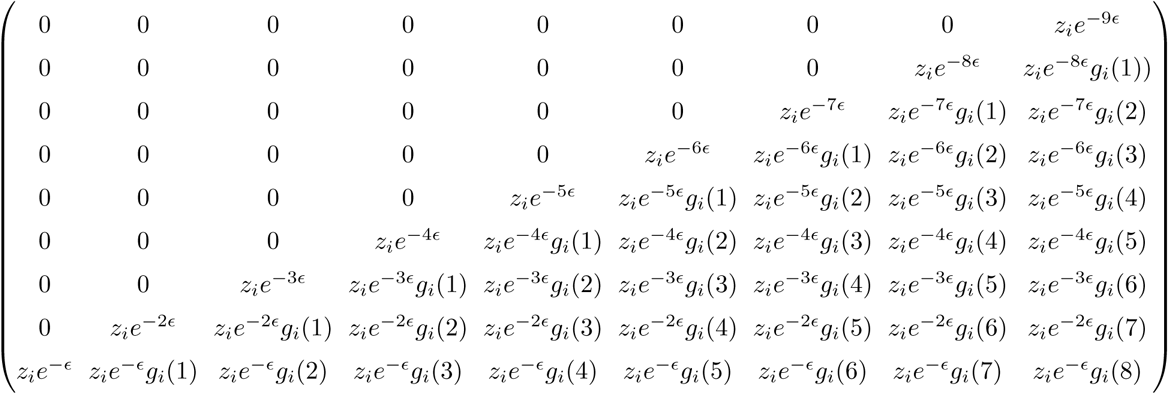

Where *z*_*i*_ is the fugacity of the attached molecule and *g*_*i*_(*n*) is the gap partition function for a gap of *n* unbound modules. Again, the matrices **M**_U_ and **M**_I_ are identical apart from the fugacity of the added molecule and the client species that binds in the gap. Since adding a SUMO molecule will leave a gap of unbound SIM modules, the gap partition function *g*_U_ describes the binding of clients composed of SUMO modules.

Applying *N* transfer matrices to the right vector generates a vector **V**(*N* + 2) describing a filament containing *N* + 2 scaffold molecules. Each element of the vector is a polynomial giving the partition function of a filament terminating with a different length sticky end. The left vectors, **V**_*i*L_, serve three purposes. First, they collapse the filament vector into a scalar polynomial that gives the complete partition function of the filament. Secondly, the left vector provides the statistical weights for client binding at the left sticky end. These two functions can be served by a vector of the form

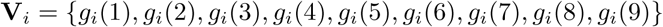

The third function of the left vector is that it must account for the fact that a filament can terminate with either species of scaffold. That means that the final left vectors are given by

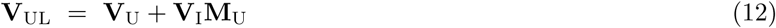

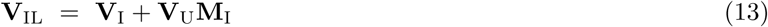

### B. Client binding entropy

The binding entropy of clients in a gap arises from the degeneracy of arranging the clients and unbound sites. We can treat the clients and vacancies as two particle types. When *N*_*c*_ clients of valence *v* bind to a gap of m sites, there are *N*_*v*_ = *m − vN*_*c*_ unoccupied sites. So the total number of particles is *N*_*t*_ = *N*_*c*_ + *N*_*v*_ = *m −* (*v −* 1)*N*_*c*_. The number of permutations is, therefore, the number of ways to select *N*_*v*_ particles from *N*_*t*_ positions. This degeneracy is gives the binomial coefficients appearing in Eqs. 5, 6.

### C. Monomer Concentration vs total concentration

The quantities *c*_U_ and *c*_I_ appearing in our calculations represent the concentration of scaffold molecules that have not formed intermolecular bonds. In contrast, the more experimentally accessible quantity is the total scaffold concentration *C*_tot_, which includes molecules that have formed assemblies. These quantities are related as follows

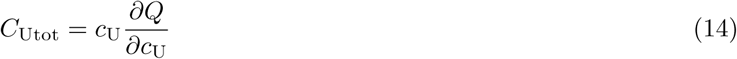

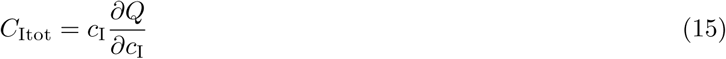

Fig 3b plots the free scaffold concentrations as a function of the total SUMO module concentration. There are several observations to make from this plot. First, when the scaffold concentrations are equal the monomer concentrations are very low, on the order of 10^*−*10^ M. This is much lower than the dilute phase concentration reported by [6]. The discrepancy is due to the fact that most molecules in the dilute phase are in the perfectly aligned dimer state. This state satisfies all the available bonding sites, rendering the scaffolds inert to further assembly processes.

Second, when one scaffold is in excess, SUMO in this case, the monomer concentrations diverge widely (note the logarithmic vertical axis). This is because the low concentration scaffolds are mostly consumed in complexes with the higher concentration module. The depletion of low concentration modules results in an excess of high concentration modules that increases as the stoichiometry asymmetry increases.

Third, it is useful to compare the concentrations in Fig 3b to the 10^5^ M^*−*1^ affinity for monovalent SUMO-SIM binding [6]. At equal scaffold stoichiometry, the scaffold monomer concentrations are much too low for monovalent binding to occur. This justifies our approximation of strictly 1D filament formation because most of the gaps only have one or two unbound sites. However, when the SUMO module concentration reaches 90 *µ*M (a 9:5 excess over SIM), the concentration of free SUMO module scaffolds is ≃14 *µ*M. This is comparable to the 0.1 ≃*µ*M concentration of clients, so monovalent binding cannot be neglected.

### D. Excess monomer scaffold binding at defect sites

When the scaffolds are present at unequal stoichiometries, the excess scaffold accumulates at concentrations where monovalent binding in the gaps becomes significant. At large asymmetry, the excess scaffold monomer concentration (≃1 *µ*M) is about ten times greater than the total client concentration (≃0.1 *µ*M) so that monomer scaffold binding is more favorable than client binding at defect sites. Therefore, we need to correct our 1D filament model to allow for perpendicular binding of scaffolds. As a first correction, we consider the “t” configuration illustrated in Fig. 7. This correction only allows for monovalent binding, which we expect to dominate given the small size of gaps in the filament.

**FIG. 7:**
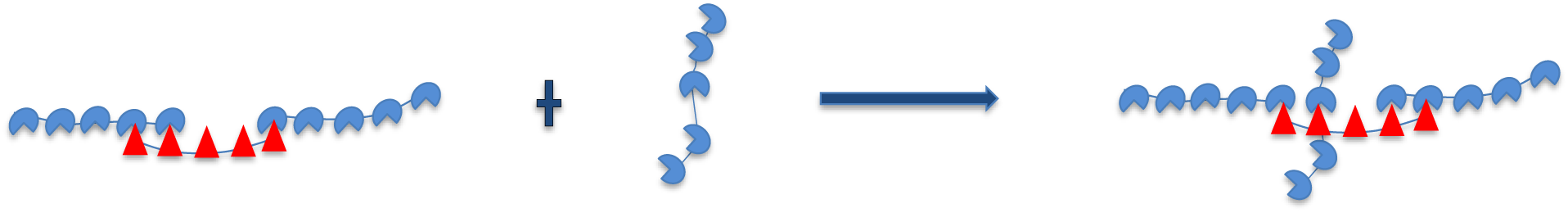
Excess monomer scaffold concentration binding at defect sites in crosslinked fashion.

The concentration of scaffolds that bind in the “t” configuration is calculated as

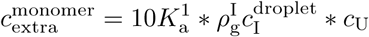

Where 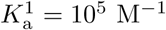 is the monovalent SUMO-SIM binding affinity [6]. This affinity is multiplied by a factor of 10 to account for the degeneracy of binding of a decavalent scaffold to a single-site defect. *c*_U_ is the scaffold monomer concentration (Figure 3b) and 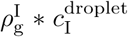 is the concentration of SIM defects in the network droplet. The latter quantity is calculated from the concentration of SIM scaffolds in the droplets, 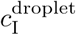 [6], and the density of defect sites per SIM scaffold, which is calculated from 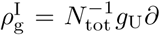ln*Q*_droplet_/∂*g*_*U*_, where we have used the substitution 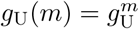 to define *g*_U_ as the statistical weight of an unbound SIM module. Again, the subscript follows from our definition of gap partition functions based on the client that they bind. Note that this correction has not been applied to our PC calculations, which explains the systematic underestimate of the PC at high SUMO concentration.

## Acknowledgments

This work was supported by NIH Grant R01GM107487 (to J.D.S), a postdoctoral fellowship from the Cancer Research Institute (to J.K.), a grant from the Welch Foundation (I-1544 to M.K.R.) and the Howard Hughes Medical Institute.

